# Depth-Resolved Macroscopic Fluorescence Lifetime Imaging via High-Spatial-Frequency Structured Illumination

**DOI:** 10.1101/2025.06.10.658928

**Authors:** Nanxue Yuan, Saif Ragab, Navid Nizam, Vikas Pandey, Amit Verma, Tynan Young, John Williams, Margarida Barroso, Xavier Intes

## Abstract

**Significance:** Macroscopic Fluorescence Lifetime Imaging (MFLI) is a powerful, non-invasive imaging modality that offers robust, physiologically relevant contrast largely independent of fluorophore concentration, excitation intensity, and tissue signal attenuation. However, accurately determining the depth of signal origin remains challenging, potentially leading to ambiguity in biological interpretation. Here, we present a novel optical correction method that effectively eliminates surface signal bias, such as that from skin in preclinical imaging, without the need for chemical clearance. This advancement supports the robust applicability of MFLI in translational research.

**Aim:** Establishment of a High Spatial Frequency–Fluorescence Lifetime Imaging (HSF-FLI) framework to selectively isolate subsurface fluorescence (deeper signals) from surface fluorescence, while preserving the accuracy of lifetime estimation.

**Approach:** A modulation transfer function (MTF) that relates spatial frequency to penetration depth was derived using Monte Carlo eXtreme (MCX) simulations (for physics-based modeling) and validated with agar-based capillary phantoms on a time-gated ICCD–DMD system. Depth-independent fluorescence was decomposed into surface and subsurface components through structured three-phase sinusoidal illumination, and nonlinear least squares fitting was applied to recover lifetime or lifetime based parameters maps. HSF-FLI was demonstrated in vivo in mouse models bearing tumor xenogratfs and was cross validated with ex vivo measurements.

**Results:** We extensively characterized the performance of High Spatial Frequency–Fluorescence Lifetime Imaging (HSF-FLI) through simulations and tissue-mimicking phantoms. The approach was further validated in vivo by assessing drug delivery in preclinical models using MFLI-FRET (Förster Resonance Energy Transfer).

**Conclusion:** By coupling structured illumination with physics-based depth modeling, HSF-FLI delivers accurate, depth-selective lifetime readouts, setting the stage for robust and fast FLI implementation in translational studies.

## 1 Introduction

Fluorescence Lifetime Imaging (FLI) has emerged as a transformative tool in molecular biology, enabling spatially resolved, quantitative measurements of key biochemical and biophysical parameters, including pH, ion concentration, oxygen tension, redox state, and molecular interactions.^1–3^ FLI quantifies the fluorescence decay time of fluorophores following excitation, offering robust and physiologically meaningful contrast that is largely independent of fluorophore concentration, excitation intensity, and signal attenuation.^1, 4^ As a result, near infrared (NIR) FLI has gained significant traction in macroscopic imaging of biological tissues (MFLI). MFLI is used in preclinical *in vivo* imaging of small animal models to study pharmacokinetics, drug delivery, immune responses, and the tumor microenvironment. Its utility is also expanding into clinical domains, including fluorescence-guided surgery and ophthalmic diagnostics.^1, 5–8^ A particularly powerful application of FLI is its ability to quantify Förster Resonance Energy Transfer (FRET), a distance-dependent energy transfer between donor and acceptor fluorophores that reports molecular interactions at *<*10 nm resolution.^6, 9^ By monitoring the reduction in donor fluorescence lifetime, FLI-FRET enables the quantitative assessment of protein–protein interactions and conformational changes in live, intact biological tissues.^5, 10^ This capability also allows for the quantification of antibody–target engagement within native tissue environments, an essential parameter for evaluating the efficacy of targeted therapeutics.^11^

At the macroscopic scale, light propagation in biological tissues is predominantly governed by the scattering properties of the sample. As a result, fluorescence signals are collected from extended three-dimensional regions, making it challenging to resolve the depth of fluorescence signal origin without more sophisticated approaches, such as fluorescence LiDAR^12^ or full 3D fluorescence tomography.^13, 14^ In the commonly used wide-field imaging configuration, each pixel can contain a mixture of signals originating from various depths, potentially introducing significant bias in biological interpretation. To address scattering effects, optical clearing agents have been developed to homogenize refractive indices within tissues, thereby improving light penetration.^15^ However, these agents are known to quench fluorescence signals^16, 17^ and may denature tissue structures *in vivo*,^18^ altering the microenvironment of the fluorophores. These alterations can compromise fluorescence lifetime measurements and introduce uncertainty in the quantification of targeted drug delivery. An alternative strategy involves using structured illumination with striped excitation patterns to modulate the light penetration depth, enabling optical sectioning to separate in-plane from out-of-plane fluorescence signals.^19, 20^ While promising, current structured illumination models are limited in their ability to determine the depth correlation of spatial frequency and have only been applied to intensity-based imaging.

To overcome these limitations, we propose a High Spatial Frequency–Fluorescence Lifetime Imaging (HSF-FLI) framework that enhances depth-resolved lifetime accuracy in macroscopic imaging. This approach is implemented on an MFLI-ICCD system, routinely used for whole-body preclinical imaging and equipped with structured light illumination capabilities. A depth-correlated modulation transfer function (MTF) was first derived through Monte Carlo simulations and subsequently validated using experimental in silico datasets. The HSF-FLI method was further evaluated in live mouse models bearing HER2+ breast tumor xenografts and treated with FRET-labeled Trastuzumab (TZM), a monoclonal antibody used for HER2-positive breast cancer therapy. In vivo, HSF-FLI was used to retrieve fluorescence lifetime and FRET-based parameters, with quantitative validation performed using corresponding ex vivo data. The excellent agreement between in vivo and ex vivo FLI parameter estimations confirmed the accuracy and robustness of the HSF-FLI approach. Moreover, these results demonstrate HSF-FLI’s ability to effectively separate superficial (skin) signals from deeper tumor-derived fluorescence in live imaging contexts.

## 2 Methods

### 2.1 MFLI ICCD Imaging System

The MFLI system is equipped with a time-gated Intensified Charge-Coupled Device (ICCD) (Picostar HR, LaVision, GmbH, Bielefeld, Germany).^21^HSF-FLI data was acquired for 101 gates with 300 ps gate width and 80 ps gate step (8 ns of overall acquisition time). An 80 MHz tunable Ti: sapphire laser (Mai Tai HP, Spectra-Physics, CA,60 USA) with a laser period of 12.5 ns provided excitation in the NIR range (690nm – 1020nm). The laser beam was directed into a digital micro-mirrors device, DMD, (DLi 4110, Texas Instruments, TX, USA) through a multi-mode optical fiber. DMD enables the projection of controlled spatial patterns encoded over 255 level of grey. The DMD was controlled a customized Labview script which projects three pre-made 8-bit sinusoidal patterns with offset of 2*π*/3 for each selected spatial frequency. Imaging settings, including emission power, Micro-Channel Plate (MCP) voltage, camera exposure time, and hardware binning ratio, were adjusted based on the pre-formulated Look-up table (LUT) of the ICCD accordingly.^22^ Fluorescence-emission filters and neutral density filters were used for FLI and IRF capturing respectively. The IRF was captured using a white paper.

In the case of FRET, the FLI based parameter of interest are typically estimated through Non-linear Least Square Fitting, NLSF, described in Equation 1. In this approach, the fluorescence decays are modeled by:

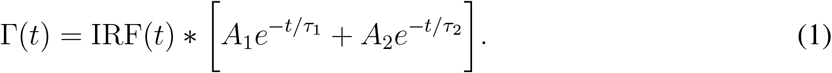

where the experimental decay, Γ(*t*), is the combination of a bi-exponential model convolved with pixel-wise instrument response function, IRF(t). The FLI based parameter of interest are the donor *τ*_1_, acceptor lifetimes *τ*_2_ and corresponding amplitude *A*_1_ & *A*_2_. The FRET percentage,*f*_1_, could thus be obtained through Equation 2.

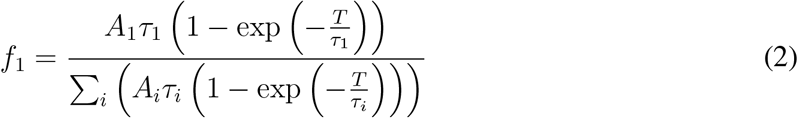

where amplitude weighted average lifetime of the two lifetime components, *τ*_*a*_, is used as the as the parameter to obtain the correct energy transfer efficiency. Equation 3.

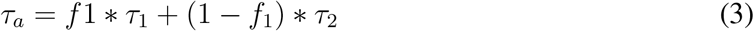

The systemic effects on the IRF and the FDC have been studied for the system earlier.^12, 22, 23^ However, the acquired fluorescence decays can be originating for different tissue depths that are not distinguishable directly from the measurements. Hence, the FLI-based parameters estimation may be affected by the heterogeneous nature of the biological tissue under investigation.

### 2.2 Spatial Frequency Domain Imaging Optical Properties Extraction

Optical properties of the agar-based 20% Intralipid phantoms are extracted through Spatial Frequency Domain Imaging, SFDI, using the MFLI system with a gate width of 1000 ps.^12^ Three pre-made 8-bit sinusoidal Labview patterns at spatial frequency of 1 *mm*^−1^ with offset of 2*π*/3 were projected through DMD. A factory-characterized silicon phantom (INO Biomimic Phantoms, QC) with known optical properties, *μ*_*a*_ = 0.03 *mm*^−1^ and 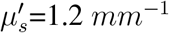, was imaged using the same imaging process to calibrate the optical properties of the sample phantom. A total of 10 gates with 500 ps step were acquired for each illumination pattern using ND filter for Intralipd and the calibration phantom under both 700 nm and 760 nm. Both phantoms were imaged at the same distance from the camera to avoid the need of performing height correction. Optical properties including *μ*_*a*_ and 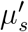 were obtained through a multi-frequency look-up table (LUT).

### 2.3 HSF-FLI MTF

HSF-FLI was derived from the optical sectioning through the deconvolution of non-depth dependent FLI signal, *I*_*DC*_, to derive the spatially modulated surface signal, *I*_*AC*_, and non-modulated subsurface signal, *I*_*sub*_. Fluorescence planar signal, *I*_*DC*_, and surface scattered signal, *I*_*AC*_, could also be demodulated based on the phase offset signals, *I*_1_, *I*_2_, &*I*_3_, due to the constant phase changing of 2*π*/3 in Equation 4.

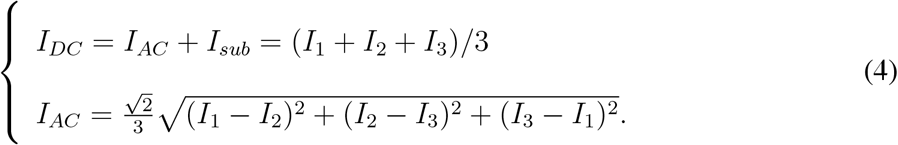

To quantify the efficiency of different spatial frequencies in HSF-FLI, Modulation Transfer Function, MTF, was derived for fluorescence imaging from MTF in Spatial Frequency Domain, SFD,^24^ and used in this paper in Equation 5,

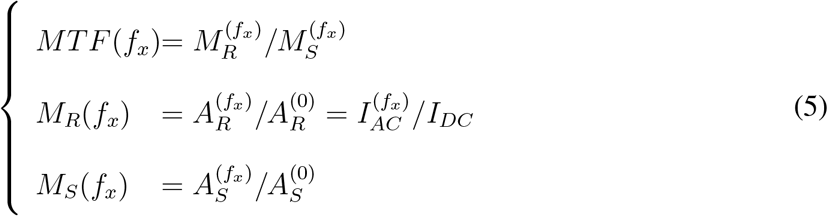

where *M*_*R*_ is the modulation depth of the reflectance from the sample and is the fraction of reflected DC amplitude, 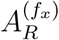, and reflected AC amplitude, 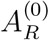, which are the same as *I*_*AC*_ & *I*_*DC*_ respectively. *M*_*S*_ is the modulation depth of source fluence and is the fraction of DC source amplitude, 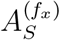, and AC amplitude, 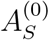, from source.

In HSF-FLI, source fluence is modulated by patterns with offsets, and the reflectance were directly captured through the Gated-ICCD detector. In this case, *M*_*S*_ could thus be calculated through the Equation 6.

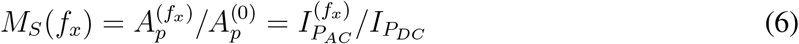

where 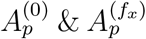 and is the amplitude of the projected patterns with DC and AC components Equation 7 and 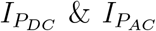 are pattern modulated source fluence of non-depth dependent and spatially modulate signal respectively.

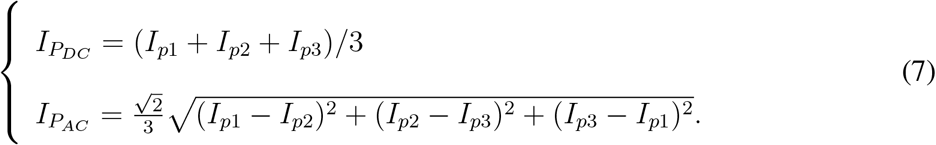

which leads to the finalized MTF equation for HSF-FLI as Equation 8

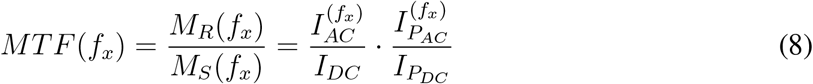

### 2.4 MCX Simulation

The details of the MCX simulation workflow to collect fluorescent emission on the surface are described elsewhere.^25^ Here, we briefly summarize the simulation workflow. The generation of fluorescence data involves a two-step process. First, fluorescent inclusions with diverse spatial and depth characteristics are embedded in computationally designed phantoms. These phantoms are illuminated using experimentally derived structured light patterns, and an MC simulation captures the resulting internal fluorescence intensities. In the second step, these internal intensities act as emission sources in a subsequent Monte Carlo simulation, propagating outward to the phantom surface. The final simulated data, representing fluorescence collected at the phantom surface, form the basis for further processing and analysis.

### 2.5 Animal Experiments

Two 4 weeks old athymic female nude mice (CrTac: NCr-Foxn1nu; Taconic Biosciences), were injected with 200 *μ*L of 10 *×* 10^6^ HER2+ AU565 breast cancer cells (ATCC) mixed 1:1 with Cultrex BME (R&D Systems Inc, Minneapolis, MN, USA) into the right and left inguinal mammary fat pads. After 4 weeks of tumor growth, NIR labeled Meditope (MDT) TZM probes were injected retro-orbitally.20 *μ*g of MDT-conjugated TZM-AF700 probe was administered retro-orbitally in the non-FRET mouse, and for the FRET mouse, 20 *μ*g MDT-TZM-AF700 (Donor)/40 *μ*g MDT-TZM-AF750 (Acceptor) FRET pair probes were injected through retro-orbital route with a difference of 2 h between donor-labeled and acceptor-labeled drug injections. Both the MDT-non-FRET and FRET pair probes were generously provided by Dr. John C. Willams lab Department of Molecular Medicine, Beckman Research Institute of City of Hope, Duarte, CA. The HSF-FLI was performed at 48hr post injection in anesthetized (EZ-SA800 System, E-Z Anesthesia, USA) mice placed on a 37 *°*C warming pad (Rodent WarmerX2, Stoelting, IL, USA) on imaging stage. After euthanasia, the tissues including skin and fat above tumors were removed carefully to expose the tumor for ex vivo imaging. The tumors were kept within mouse body to reduce the effects of changing fluorescence environments. All animal protocols were conducted with approval by the Institutional Animal Care and Use Committee at both Albany Medical College and Rensselaer Polytechnic Institute.

## 3 Results

MCX simulations were performed on a tilted capillary phantom and directly compared with experimental measurements to assess whether the simulated datasets can be used to correlate spatial frequency with depth information in real-world applications in Figure 1. The bulk medium was assigned optical properties of *μ*_*a*_ = 0.002 mm^−1^ and 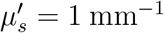, while the interior dye solution was set to *μ*_*a*_ = 0.04 mm^−1^ and 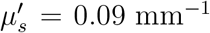. In the forward simulation, the black surface acted as the plane of illumination; in the reverse simulation, it served as the plane of detection as shown in Figure 1 (a) left. For validation, the capillary phantom was filled with 1 *μ*M Alexa Fluor 700 in an intralipid scattering medium shown in Figure 1 (a) right. Data were acquired at spatial frequencies from 0.1 to 0.7 *mm*^−1^ in 0.1 *mm*^−1^ steps; results at 0.2 and 0.6 m*mm*^−1^ are shown here, with the full frequency series in Supplementary Figs. S1 and S2.

**Fig 1.**
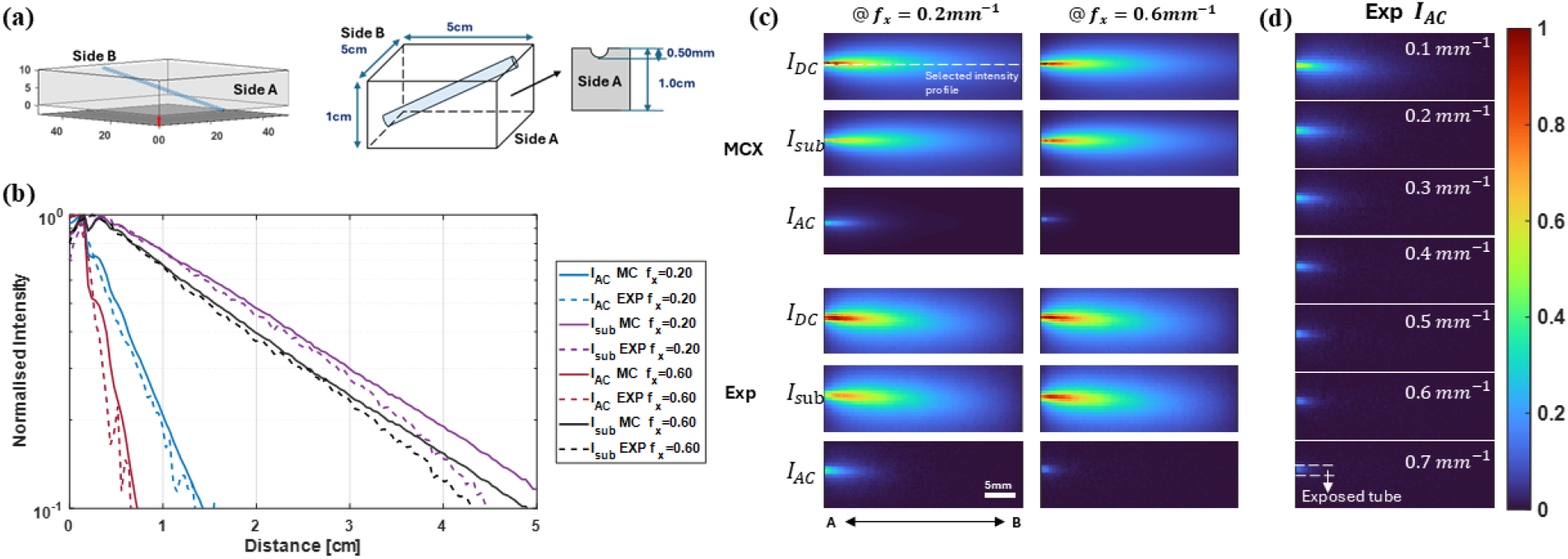
Comparison of fluorescence intensity profiles between MCX simulations and experiments, (a) the structure of MCX phantom on the left, and design of the experimental agar-based capillary phantom filled with 1 *μ*M AF700 on the right. (b) Normalized Intensity distribution from side A to B at selected fx follow selected intensity profile in dotted line. (c) Normalized fluorescence planar signal, *I*_*DC*_, and calibrated subsurface signal, *I*_*AC*_, of a tilted tube at three different spatial frequencies, fx = 0.2, 0.4, 0.6 *mm*^−1^. (d) Experimental *I*_*AC*_ changing as spatial frequency increasing

To compare simulation and experiment, the profiles of *I*_*AC*_ and *I*_*sub*_ were extracted along the A–B axis and normalized to the peak value of each profile in Figure 1 (b), and the selected intensity profile was shown in the white dotted line in Figure 1(c). A section of the tube was deliberately exposed on side A to provide a surface reference; profiles exceeding unity are displayed. Despite the noise in the experimental curves, both *I*_*AC*_ and *I*_*sub*_ agree closely between simulation and measurement at the selected frequencies and the corresponding values of *R*^2^ are included in the Supplementary Information Table. S1. Finally, after normalizing by *I*_*DC*_, we compared *I*_*DC*_, *I*_*AC*_, and *I*_*sub*_ across all frequencies: *I*_*sub*_ amplitude increases with spatial frequency—indicating shallower sampling depths—whereas *I*_*AC*_ captures deeper regions at low frequencies and shifts toward shallower depths as the frequency increases in Figure 1 (d).

Under high spatial-frequency conditions, accurate pattern resolution is critical for analysis. Given the limitations of the gated ICCD and DMD system, it is equally important to determine the maximum spatial frequency that can be achieved without information loss. In Supplementary Fig. S2, the coefficient of determination (*R*^2^) for *I*_*AC*_ decreases from 0.957 to 0.926 as the spatial frequency increases from 0.6 to 0.7 mm^−1^, while *I*_*sub*_ remains approximately constant. These results indicate that sinusoidal patterns above 0.6 mm^−1^ cannot preserve projection accuracy; therefore, the system’s maximum usable spatial frequency is 0.6 mm^−1^.

With the validity of MCX simulations established for mimicking real in silico conditions, we quantified how fluorescence-signal sensitivity varies with spatial frequency via the modulation transfer function (MTF) defined in Equation 8. Depth-resolved MTF profiles—spanning from 0.50 mm to 9.50 mm into the capillary—are presented in Figure 2 (a) and the simulation schematic are shown in Figure 2 (b). Spatial frequency was incremented in 0.01 *mm*^−1^ steps. At the zero-frequency limit (*f*_*x*_ = 0*mm*^−1^), *I*_*AC*_ and *I*_*DC*_ are identical, yielding an MTF of 1. As frequency increases, the MTF steadily declines across all depths, reflecting reduced ability to resolve finer patterns at greater penetration. At selected spatial frequency for experimental conditions, *f*_*x*_ = 0.6*mm*^−1^, the MTF drops below 0.01, two orders of magnitude down, beyond depths of roughly 1.5 mm to 1.75 mm, indicating that this frequency represents the practical resolution limit in depth. This trade-off between spatial resolution and penetration depth underscores the importance of selecting frequencies that balance pattern fidelity with sufficient sampling depth in MFLI experiments.

**Fig 2.**
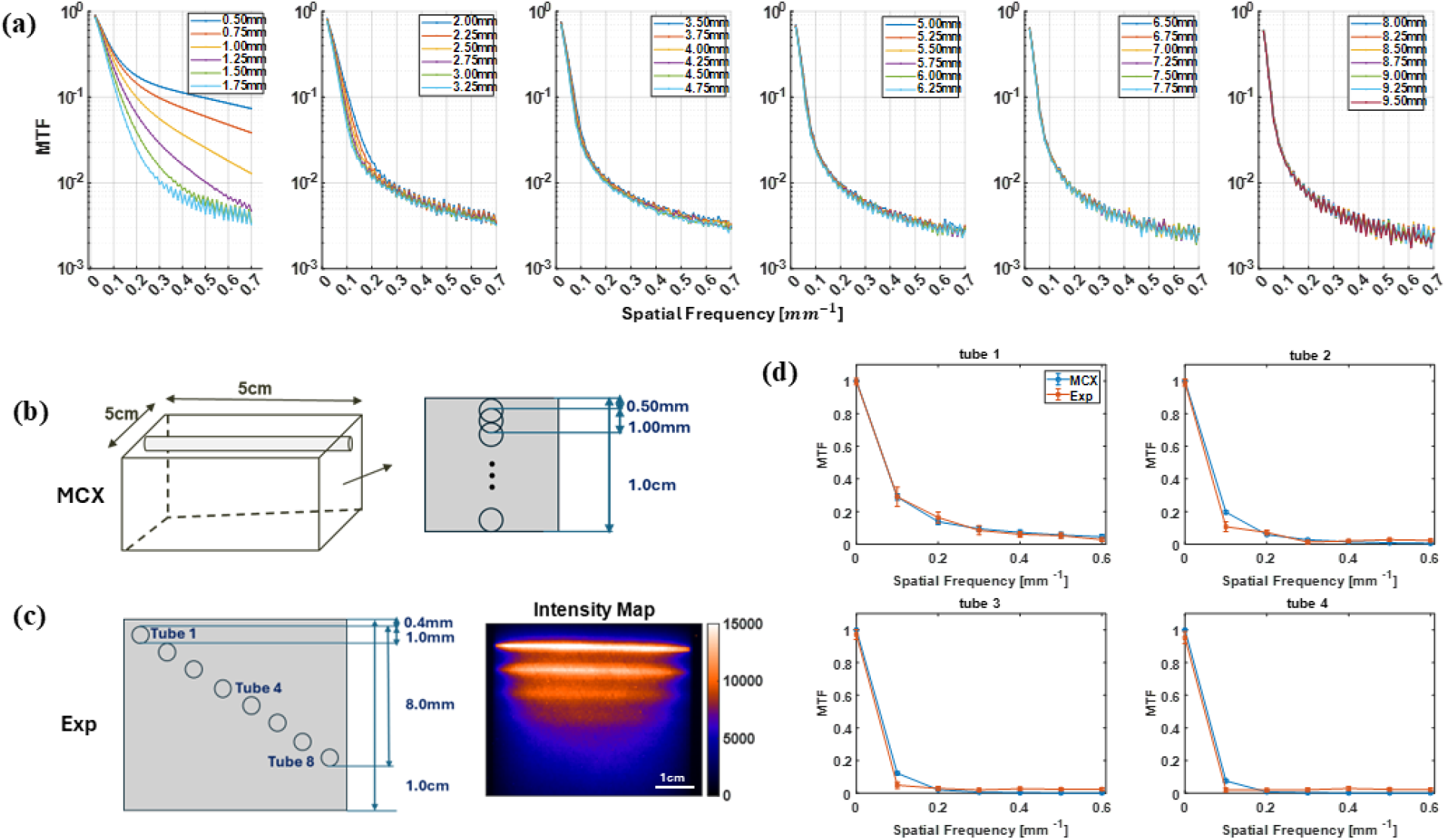
Variation of MTF with spatial frequencies, (a) depth-related MTF through MCX simulated fluorescence data, fx = 0.01 to 0.70 *mm*^−1^ in steps of 0.01 *mm*^−1^. (b) MCX phantom design with capillaries shifted downwards progressively in steps of 0.25 mm. (c) Experimental agar-based capillary phantom, 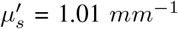 with 700nm excitation wavelength through SFDI, designed with 8 1mm glass capillaries filled with 1 *μ*M AF700 sample. Tube 1 is approximately 0.4mm beneath the surface, and the corresponding intensity Map. (d) MTF comparison between first 4 tubes in experimental phantom and MCX at the same depth with fx = 0.0 to 0.60 *mm*^−1^ in steps of 0.01 *mm*^−1^.

To validate the accuracy of the MCX generated MTF, the experimental phantom was designed to have 8 1mm tubes in total in agar-based intralipid phantom with *μ*_*a*_ = 0.02*mm*^−1^ & 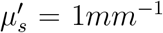 characterized under 700nm excitation wavelength. The structure of the phantom is shown in Figure 2(c). Each tube was filled with 1 *μ*M AF700, producing the fluorescence intensity map shown in Figure 2(c) right. In the phantom experiment, only first 4 tubes were detectable in the intensity map and they were further selected as the targets for comparing the MTF with MCX simulated MTF at similar heights shown in Figure 2(d) at depths of 0.4 mm, 1.4 mm, 2.4 mm, and 3.4 m. Shallower tubes exhibited sharper boundaries due to reduced scattering, while deeper tubes appeared progressively diffuse. At 0.4 mm depth, simulated and experimental MTF curves aligned with an R^2^ of 0.98 across spatial frequencies from 0 to 0.6 *mm*^−1^. Correlation coefficients decreased with depth (*R*^2^ = 0.95 at 1.4 mm, 0.91 at 2.4 mm, and 0.87 at 3.4 mm), reflecting increased scattering and signal attenuation. From the comparison between both the Intensity profile shown in Figure 1 and MTF of MCX and experimental results, the MCX simluation showed good correspondce as the experiemtnal, and the MCX-derived MTFs accurately emulate real fluorescence-lifetime measurements and are suitable for direct application in experimental settings.

Following the strong agreement between MTFs from MCX simulations and phantom measurements, HSF–FLI was implemented in the time domain to assess fluorescence-lifetime accuracy after removal of surface-scattering contributions, *I*_*AC*_. Analysis was limited to the three shallowest tubes of the capillary phantom depicted in Figure 2. Figure 3(a) presents fluorescence-decay curves, FDCs, of the depth-independent signal *I*_*DC*_ from these tubes alongside a 1 *μ*M AF700 reference acquired under scattering-free conditions in a well plate. All images for phantom and wellplate were captured using identical instrument settings to avoid lifetime-analysis bias from varying noise conditions. In the absence of scattering, the AF700 well-plate signal peaked at approximately 2300. Under scattering in tube 1, the peak intensity decreased by roughly 50 %, with further reductions observed at greater tube depths.

**Fig 3.**
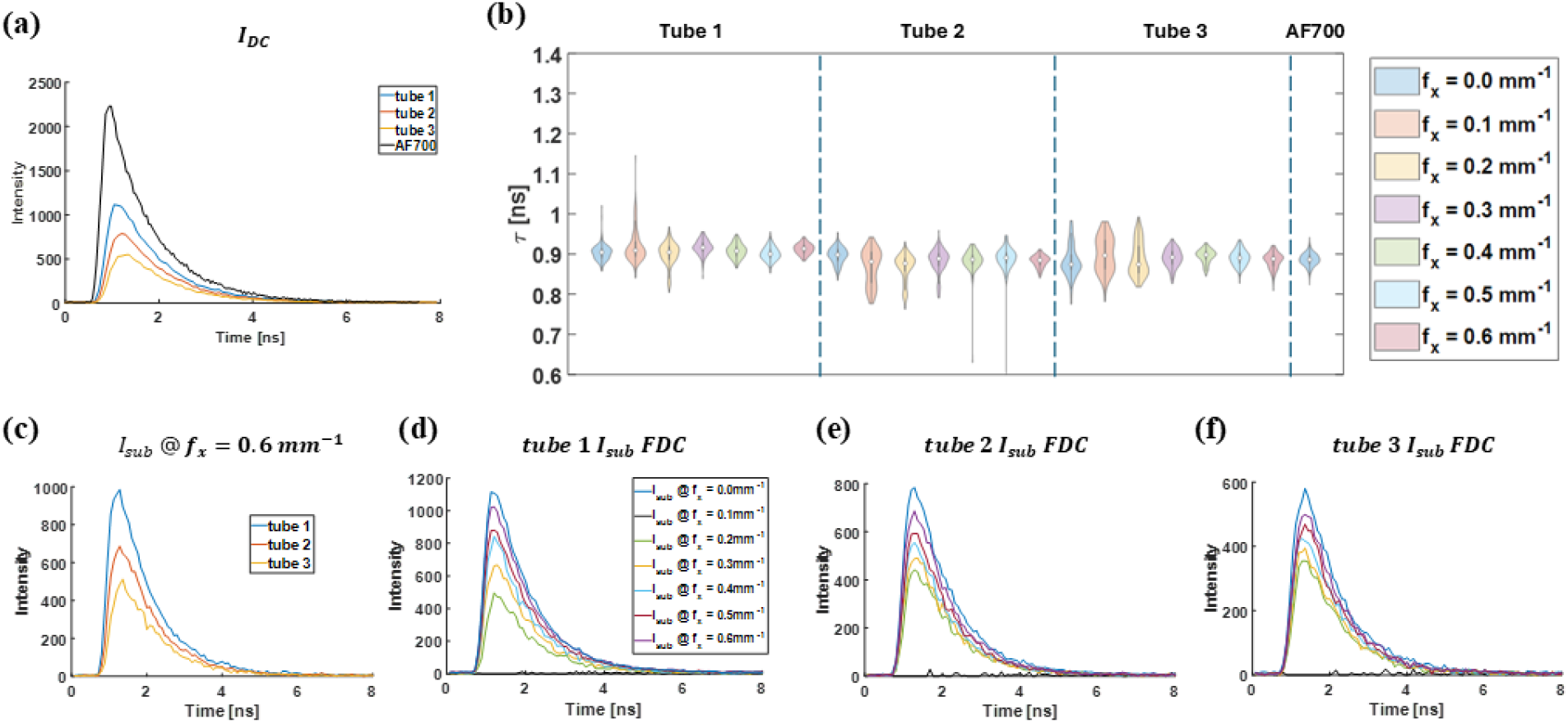
Time domain HSF-FLI of first three tube in Figure 2, (a) Fluorescence Decay Curve (FDC) of *I*_*DC*_, for three tubes and the control fluorescence dye in wellplate, 1 *μ*M AF700. (b)lifetime results from NLSF fitted *I*_*sub*_ at different spatial frequencies of tube 1, 2, & 3, and wellplate AF700 sample.(c) Reconstructed *I*_*sub*_ at fx = 0.6 *mm*^−1^ comparison among three tubes. Reconstructed *I*_*sub*_ for fx = 0.10 to 0.60 *mm*^−1^ in steps of 0.01 *mm*^−1^ of (d) tube 1, (e) tube 2, (f) tube 3.

Following removal of the surface signal, *I*_*sub*_ at each spatial frequency underwent non-linear least-squares fitting (NLSF) for each of the three capillary tubes; the resulting lifetimes are displayed in Figure 3(b). Standard deviation of the fitted lifetimes declined as spatial frequency increased, while the mean lifetime remained near 0.9 ns across all tubes. Deeper tubes exhibited larger overall lifetime variability, correlating with reduced fluorescence-decay signal quality shown in Figure 3(d–f). Representative fluorescence-decay curves for *I*_*sub*_ at 0.6 *mm*^−1^ shown in Fig. 3 (c) reveal depth-dependent noise artifacts that drive NLSF uncertainty, as reflected in the standard deviations plotted in Figure 3(b). At approximately 0.4 mm depth (tube 1), lifetime distributions for the capillary and the scattering-free AF700 reference converge at 0.4 *mm*^−1^ and exhibit the smallest spread at 0.6 *mm*^−1^. At 1.4 mm (tube 2), standard deviations increase for all spatial frequencies but remain comparable to those of the well-plate sample. In the deepest tube (2.4 mm), standard deviations continue to decrease with higher spatial frequencies, although slight fluctuations in mean lifetime likely arise from the markedly lower signal intensity.

The HSF-FLI was further applied to non-FRET and FRET mouse models bearing AU565 tumor xenografts. *In vivo* and *ex vivo* imaging was performed 48hr post-injection of NHS-conjugated MDT-TZM-AF700 non-FRET or MDT-TZM-AF700/MDT-TZM-AF750 FRET pair probes shown in Figure 4 and Figure 5. Traditional NHS-ester dye labeling leads to random, non-specific conjugation that compromises reproducibility, functional integrity, and quantitative accuracy. To overcome these limitations, we have used MDT-TZM, an engineered antibody allowing site-specific labeling via a 12-amino acid cyclic peptide binding site in the Fab region. This modular system enables consistent dye conjugation without affecting HER2 binding or pharmacokinetics, improving imaging reliability, delivery, and therapeutic potential.^26^ Therefore, in the present study we have used MDT-TZM conjugated with donor and acceptor NIR dyes as FRET probes to enable precise, reproducible, and quantitative imaging of HER2-positive tumors.

**Fig 4.**
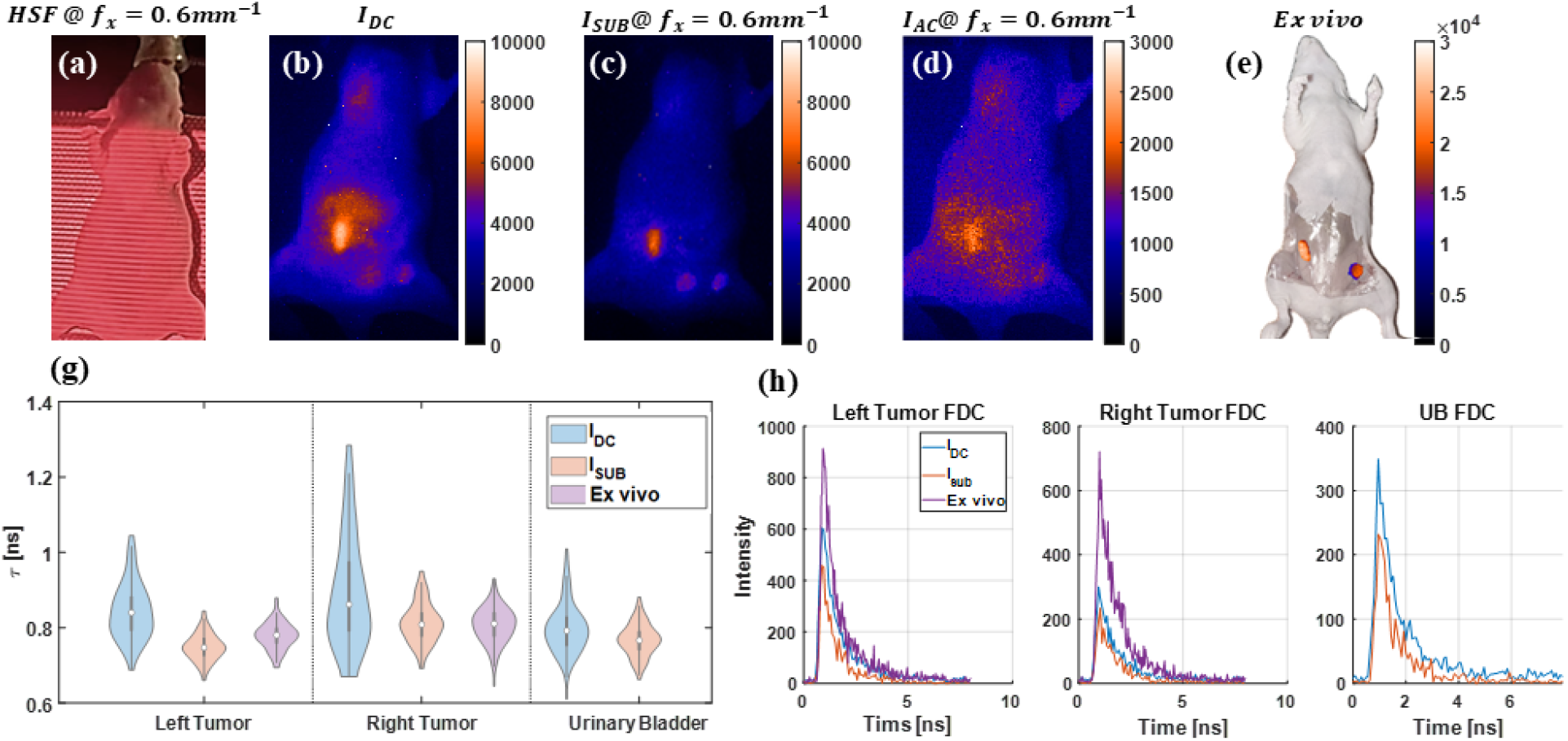
Time domain HSF-FLI of in vivo sample injected with MDT-TZM-AF700, (a) HSF projection of in vivo sample at fx = 0.6 *mm*^−1^, Fluorescence intensity map of in vivo sample at fx = 0.6*mm*^−1^ of (b)*I*_*DC*_, (c) *I*_*sub*_, (d) *I*_*AC*_. (e) Fluorescence intensity map of skin opened ex vivo sample, tissue above tumors removed after euthanasia, skin removed ex vivo mouse. (f) Lifetime comparison of left tumor, right tumor, and urinary bladder from NLSF fitting of *I*_*DC*_, *I*_*sub*_, and *ex vivo* skin opened mouse FDCs. (g) FDCs used for NLSF fitting.

**Fig 5.**
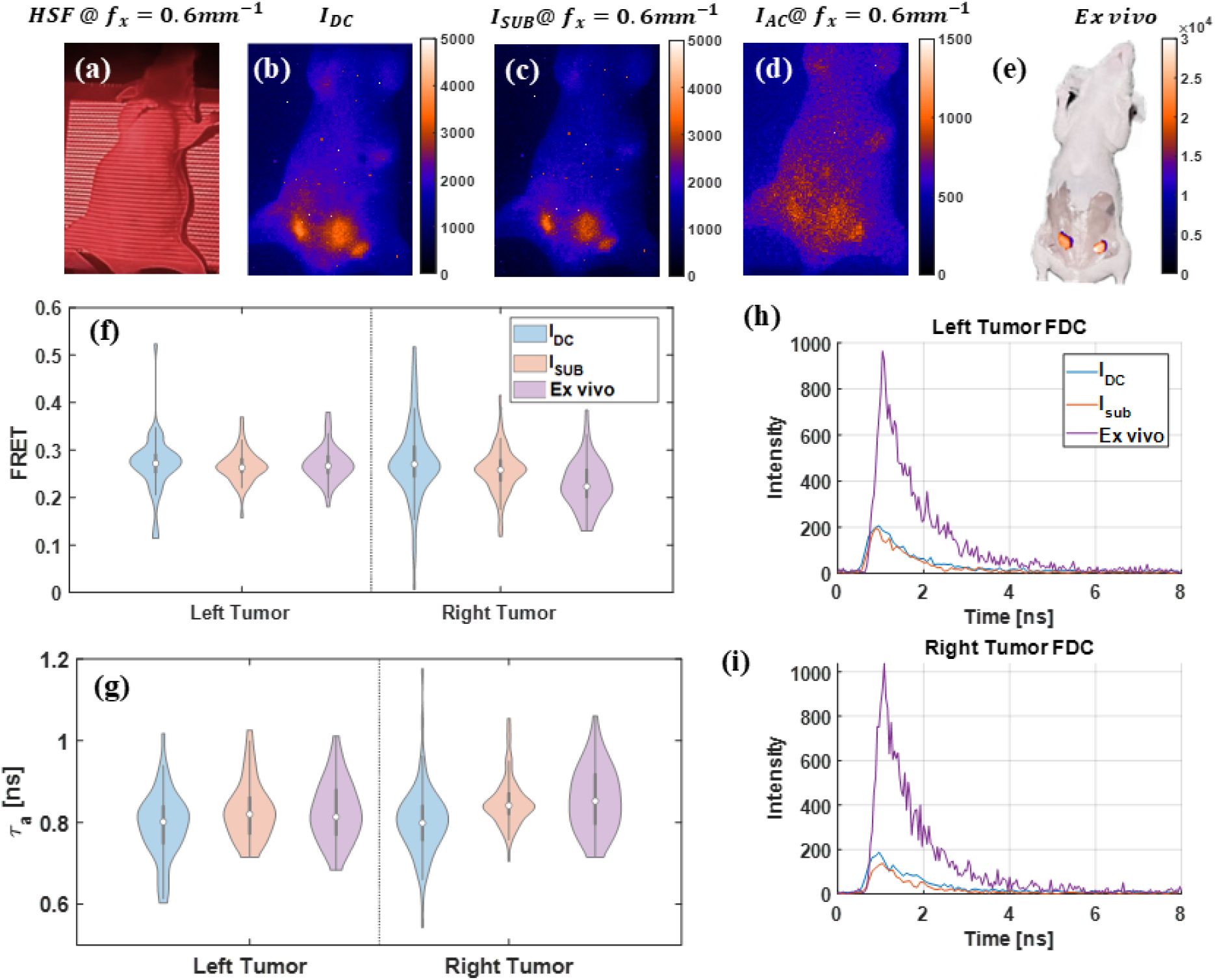
Time domain HSF-FLI of in vivo sample injected with MDT-TZM-AF700/MDT-TZM-AF750 FRET pair, (a) HSF projection of in vivo sample at fx = 0.6 *mm*^−1^, Fluorescence intensity map of in vivo sample at fx = 0.6*mm*^−1^ of (b)*I*_*DC*_, (c) *I*_*sub*_, (d) *I*_*AC*_. (e) Fluorescence intensity map of skin opened ex vivo sample, tissue above tumors removed after euthanasia. Comparison of left tumor and right tumor, and urinary bladder from NLSF fitting of *I*_*DC*_, *I*_*sub*_, and *ex vivo* skin opened mouse FDCs for (f)FRET, and (g)Amplitude weighted mean lifetime, *τ*_*a*_, (h) FDCs used for NLSF fitting.

Based on reported female mouse skin thicknesses of 400–500 *μ*m,^27^ a spatial frequency of 0.6 *mm*^−1^ was chosen, and the structured-light pattern on the *in vivo* sample is shown in Figure 4(a). In the wide-field, non-depth-resolved fluorescence image *I*_*DC*_, tumor boundaries appear indistinct, and high signal from the gut overlaps with the left tumor shown in 4(b). Excess fluorescence from non-tumor regions was eliminated by HSF–FLI, yielding clear delineation of both tumors and the urinary bladder once surface signal *I*_*AC*_ is removed in 4(c),(d). Upon removal of tissue above tumor region, including skin and fat, signal intensity increases approximately threefold in 4(e), creating a scattering-free reference. Lifetime maps for tumors and bladder are presented in Figure 4(f). Under scattering conditions *I*_*DC*_, lifetime distributions across all three ROIs exhibit broad variability; in contrast, both surface-signal-subtracted *I*_*sub*_ and *ex vivo* measurements show similar distributions and standard deviations. The absence of a urinary bladder signal in ex vivo imaging is attributed to post-mortem urine expulsion, and thus was excluded from lifetime analysis. High MCP voltage imaging settings in both *in vivo* and *ex vivo* experiments introduce noise into FDCs in Fig. 4(g), although peak intensities remain comparable to phantom and fluorophore measurements in Fig. 3. Despite these limitations, HSF–FLI reliably enhances ROI boundary contrast and suppresses superficial scattering, thereby improving fluorescence-lifetime accuracy.

HSF–FLI was applied to the FRET mouse bearing staggered injections (2hr interval between donor and acceptor injection) of MDT-TZM-AF700 and MDT-TZM-AF750, with the structured-light pattern, 0.6 *mm*^−1^, generated by the DMD shown in Figure 5(a). In the wide-field, non-depth-resolved intensity map *I*_*DC*_, fluorescence signal from the lower abdomen spreads across tumor and bladder ROIs, rendering their boundaries indistinct; additional signal spillover from the liver and upper-left flank is also apparent in Figure 5(b). Under high-spatial-frequency illumination, undesired superficial contributions are effectively suppressed, producing separable ROIs in Figure 5(c,d). Following euthanasia and superficial tissue removal, the intensity map shown in Figure 5(e) exhibits a sixfold increase in peak signal relative to the in vivo condition. FRET efficiencies were quantified via bi-exponential NLSF according to Equation 2, and the resulting standard deviations for both tumors decreased with high-frequency illumination and tissue removal in Figure 5(f). Amplitude-weighted lifetimes,*τ*_*a*_ calculated as Equation 3, which reflect donor quenching, follow the same trend as Figure 5(g). Fluorescence-decay curves used for FRET fitting are presented in Figure 5(h,i); although overall in vivo intensities decrease due to donor quenching, decay profiles remain noisy owing to the high-gain imaging settings.

## 4 Discussion

In summary, the HSF-FLI framework leverages structured illumination to isolate subsurface fluorescence signals and thereby improve the accuracy of lifetime measurements in scattering media. It translates structured-illumination optical sectioning into the time-domain by exploiting the depth-dependent attenuation of spatial frequencies. By deriving an HSF-FLI modulation transfer function (MTF) through both MCX simulations and experimental phantoms, a strong correlation (with *R*^2^ *≥* 0.87 across depths up to 3.4 mm) between simulated and measured data. This robust agreement confirms that the MCX simulation pipeline can accurately predict the depth-dependent attenuation of high-frequency structured light in turbid media, and thus provides a essential quantitative basis for selecting spatial frequencies that suppress surface-scattered light while preserving subsurface fluorescence in MFLI system.

Depth-independent fluorescence *I*_*DC*_ was de-convoluted into surface-scattered *I*_*AC*_ and subsur-face *I*_*sub*_ components via phase-shifted sinusoidal illumination from DMD. Nonlinear least-squares fitting of *I*_*sub*_ revealed a systematic reduction in lifetime variability as spatial frequency increased, the standard deviation of fitted lifetimes declined by nearly 50% at 0.6 *mm*^−1^, while the mean lifetimes remained within 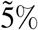 of scattering-free references. These improvements were most pronounced in the shallowest phantom tube (0.4 mm depth), but remained significant even at depths of 4.4 mm, where lifetime precision under wide-field illumination was otherwise compromised by severe photon scattering.

The application of *in vivo* and *ex vivo* to both non-FRET and FRET mouse models further confirmed the efficacy of the method by consistently suppressing superficial fluorescence contributions and improving the delineation of tumors and the urinary bladder. In non-FRET experiments, life-time distributions from *I*_*sub*_ closely matched *ex vivo* measurements after skin removal, confirming that the HSF-FLI method can effectively negate the confounding effects of tissue scattering. In FRET studies, high spatial frequency illumination reduced the standard deviation of calculated FRET efficiencies in both tumors compared to wide-field data, despite increased noise from high MCP gain imaging settings. These findings underscore HSF-FLI’s capability to enhance quantitative accuracy in both lifetime and FRET measurements under realistic biological conditions.

Despite these promising findings, several limitations merit consideration. First, the practical maximum spatial frequency in the current ICCD–DMD system is constrained to approximately 0.6 *mm*^−1^; beyond this threshold, the coefficient of determination for *I*_*AC*_ drops from 0.957 to 0.926, indicating substantial pattern distortion. Future hardware improvements, such as higher-resolution DMD chips, faster gating electronics, or adaptive optics, could extend this limit and enable finer depth discrimination. Additionally, higher spatial frequencies inherently reduce photon throughput, leading to lower signal-to-noise ratios at greater depths. Mitigation of these trade-offs may involve optimizing illumination power, detector settings, or complementary photon-collection strategies. Overall, HSF-FLI provides a robust, experimentally validated pathway to mitigate tissue-scattering artifacts in MFLI, enabling more reliable *in vivo* molecular investigations.

Future extensions of the HSF-FLI paradigm include combining multiple spatial frequencies in a single acquisition to reconstruct depth-resolved lifetime maps, integration with advanced lifetime-analysis techniques, phasor plots or global fitting, to enhance noise robustness and fasten the data processing. Moreover, translating HSF-FLI into the short-wave infrared (SWIR) window may of-fer deeper penetration and lower background auto-fluorescence, particularly valuable for clinical intra-operative imaging. Prospective applications in guided drug-delivery and receptor-ligand engagement studies—particularly those employing FRET-based reporters—stand to benefit from the improved quantitative accuracy and spatial specificity demonstrated here.

## 5 Conclusion

In this study, the HSF-FLI framework enabled depth-selective macroscopic lifetime measurements by exploiting the differential attenuation of structured illumination in scattering media. Close agreement between MCX simulations and phantom experiments validates its ability to suppress surface fluorescence and isolate subsurface signals, markedly improving lifetime precision while preserving true decay values. *In vivo* and *ex vivo* studies confirm sharply defined lifetime maps and enhanced accuracy in FRET assays, despite realistic noise levels. Although pattern fidelity and photon throughput diminish at very high spatial frequencies, these challenges can be addressed through optimized illumination and detection. HSF-FLI thus offers a robust framework for high-precision molecular imaging in living tissues.

## Code, Data, and Materials Availability

All data and MATLAB routines are available upon request.

## Acknowledgments

This work was funded by the National Institute of Health (NIH) under grants R01CA271371, R01CA237267, R01CA207725 and R01CA250636.

## First Author

Biographies and photographs of the other authors are not available.

