## Supplementary Information for "Depth-Resolved Macroscopic Fluorescence Lifetime Imaging via High-Spatial-Frequency Structured Illumination"

**Nanxue Yuan[a,*], Saif Ragab[a], Navid Nizam[a], Vikas Pandey[b], Amit Verma[c], Margarida Barroso[c], Tynan Young[d], John Williams[d], Xavier Intes[a,b]**
[a]Department of Biomedical Engineering, Rensselaer Polytechnic Institute, Troy, New York 12180, USA
[b]Center for modeling, simulation and Imaging in Medicine, Rensselaer Polytechnic Institute, Troy, New York 12180, USA
[c]Department of Molecular and Cellular Physiology, Albany Medical College, Albany, New York 12208, USA
[d]City of Hope, Beck Research Institute, Department of Molecular Medicine, Duarte, CA 91016, USA

### 1 Introduction

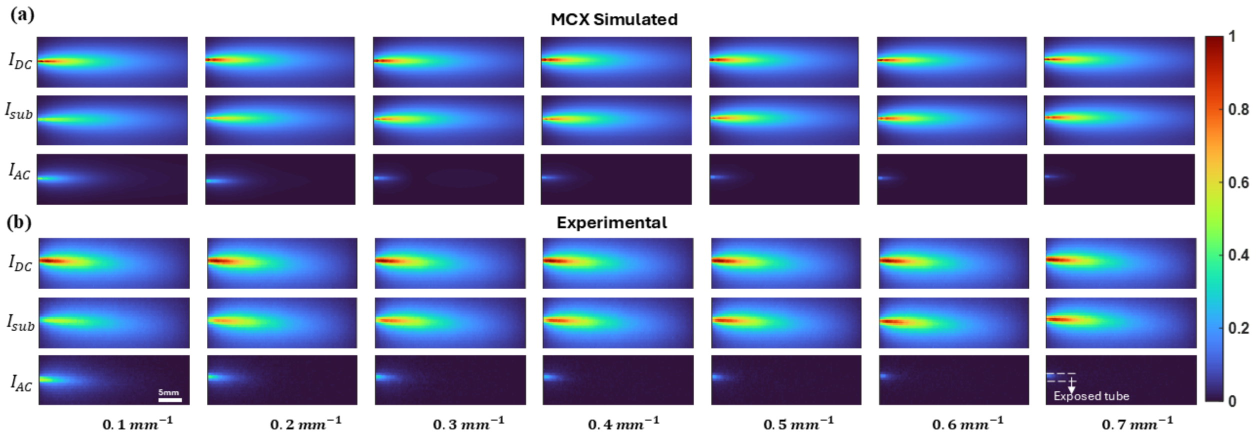


**Fig 1** Comparison of fluorescence intensity profiles between MCX simulation (a) and experiments (b). Normalized fluorescence planar signal, $I_{DC}$, and calibrated subsurface signal,$I_{AC}$, of a tilted tube at fx = 0.1 to 0.7$mm^{-1}$ in steps of 0.1 $mm^{-1}$.

| | Spatial frequency [$\mathrm{mm}^{-1}$] | | | | | | |
|---|---|---|---|---|---|---|---|
| | 0.10 | 0.20 | 0.30 | 0.40 | 0.50 | 0.60 | 0.70 |
| $I_{\mathrm{AC}}$ | 0.993 | 0.988 | 0.975 | 0.953 | 0.970 | 0.957 | 0.926 |
| $I_{\mathrm{sub}}$ | 0.972 | 0.982 | 0.988 | 0.987 | 0.989 | 0.990 | 0.989 |

**Table 1** $R^2$ of MXC simulated and Experimental intensity differences at spatial frequency range from 0.1 to 0.7 $mm^{-1}$ by 0.1 $mm^{-1}$.

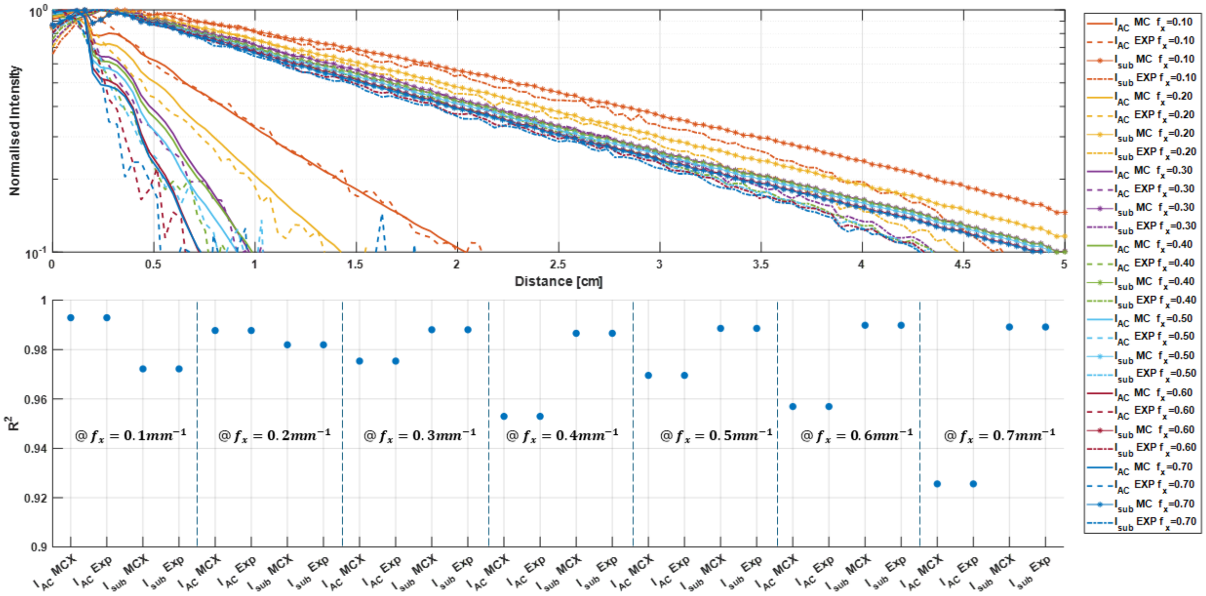

**Fig 2** Comparison of normalized Intensity distribution from side A to B follow selected intensity profile in dotted line at fx = 0.1 to 0.7$mm^{-1}$ in steps of 0.1 $mm^{-1}$ for (a) MCX simulated and experiment of $I_{AC}$ and $I_{sub}$ separately, (b) corresponding $R^2$.
